# A first-principle for the nervous system

**DOI:** 10.1101/087353

**Authors:** Kunjumon I. Vadakkan

## Abstract

Higher brain functions such as perception and memory are first-person internal sensations whose mechanisms can have options to concurrently activate motor neurons for behavioral action. By setting up all the required constraints using available information from different levels, a theoretical examination from a first-person frame of reference led to the derivation of a first-principle of the structure-function units. These units operate in synchrony with the synaptically-connected neural circuitry. These units are capable of explaining and interconnecting findings from various levels and enable triangulation of a large number of observations from both the normal and “loss of function” states of the system. Indirect evidence for the presence of a comparable circuitry in a remote species − *Drosophila* − suggests the universal presence of these units across the animal kingdom. The key features of the basic unit meet the expectations of the K-lines proposed for the development of artificial intelligence.

## 1 Introduction

Learned-information is retrieved in the form of first-person internal sensations. The lack of third-person sensory experience for the internal sensations of different higher brain functions limits empirical research. In this context, by using a large number of third-person observations from various levels, it is necessary to derive a theoretically-fitting first-principle from which all the nervous system functions can be explained. It is also necessary to understand its structural details that can provide unique circuit properties capable of interconnecting data collected from various levels (Sejnowski et al., 2014). At a minimum, it should be a testable mechanism that can operate in synchromy with the synaptically-connected neural circuitry, able to explain a mechanism for the induction of units of internal sensations occurring concurrently with behavioral motor action, suitable to explain different higher brain functions from the modifications of the above mechanism, and able to explain different nervous system disorders as “loss of function” states of the proposed mechanism. This can be further tested for interconnected explanations for alarge number of features of the system (Abbott, 2008; Edelman, 2012) (Supplementary Information I), which implies that the solution is a unique one.

The steps towards deriving the solution have to undergo different cycles of testing and retesting to adjust the backbone of the principle axiom that can lead to the arrival of a deep-rooted first-principle such that all the other findings in the system can make sense with respect to it. Even though it required working in the absence of direct empirical evidence at the time of its proposition, the present work explains the logical steps that were necessary to arrive at a solution. This approach used procedures similar to that of finding a solution for a set of algebraic equations (Supplementary Information II). To arrive at the correct solution, it is necessary to include all the non-redundant functional features of the system. Ignoring one or few non-redundant properties can result in arriving at a wrong solution. This is explained through an example of a system of equations in multiple variables (Supplementary Information III). The solution was further verified by the method of triangulation using the findings from different levels in both normal and “loss of function” states of the system.

## 2 Derivation of the mechanism

### 2.1 Constraints to work with

It is necessary to work with the following constraints to arrive at a solution. There should be a specific signature for each associative learning-induced change from which internal sensations of memory can be induced. Related associative learning events should lead to an unambiguous clustering of these associations to form “islets.” There should be specific coding for each association within each islet so that they can be specifically activated. Each new associative learning event will add new signatures pertaining to the unique features of the sensory stimuli. Continuation of this leads to both an extension of the association code and their self-organization. Since several physical properties of the items and stimuli from events in the environment are common, a) several of the previously formed islets of associations will be shared by learning events, and b) the internal sensations induced by the shared properties of the items and events form a background matrix upon which specific internal sensory units form specific memory for efficient operations. There should be a robust method to keep the unrelated islets separate from each other. Any defect in maintaining the specificity of associations within an islet will lead to false conjunctions that will be expressed as “loss of function” states of the system. These may manifest either as hallucinations, loss of internal sensation of memories in response to specific cue stimuli or defects in other higher brain functions and motor responses.

### 2.2 Derivation

First, it is necessary to identify a brain function that has a fair sample of third-person observed changes at multiple levels and that can be manipulated by experiments. In this regard, learning and memory have the advantages that multiple associative learning events can be carried out, memory can be retrieved by exposure to specific cue stimuli, surrogate behavioral motor actions can be studied and artificial changes can be made at different levels. The qualia of the internal sensations of working, short- and long-term memories are almost similar in nature. A lack of cellular changes during memory retrieval indicates that the internal sensation of memory results from passive reactivation of learning-induced changes. This is expected to take place from reactivatible (for memory retrieval), augmentable (motivation-promoted learning), reversible (forgetting), and stabilizable (long-term memory) cellular changes at the time of learning. Since related-learning is more efficient than novel learning, it occurs by sharing certain previous learning-induced changes (Tse et al., 2007). This indicates the necessity of retrieving elements of memories using changes induced by different learning events. Such an operation is expected to be efficient if each learning event consists of several unitary changes that can be used later by the sensory components of any cue stimulus. The cue stimulus is expected to induce units of internal sensations that undergo a computational process at physiological time-scales. For example, rapidly changing a general cue stimulus step-by-step towards a specific one leads to corresponding changes in the retrieved memories from a general to a specific one. The cue-directed rapid induction of changing memory is expected to result from the natural computation of several units of internal sensations.

During associative learning of two stimuli, the inputs are expected to converge at certain locations to induce a specific change such that at a later time, the presence of one stimulus can reactivate this change for inducing the internal sensation of memory of the second stimulus (Fig.1a). The next step is to find out the correct level at which the convergence of the associatively-learned stimuli can occur. Each neuron in the visual and motor cortices of a monkey has nearly 5.6 × 10^3^ to 60 × 10^3^ dendritic spines (spines or postsynaptic terminals) (Cragg, 1967). Postsynaptic potentials from nearly forty spines can spatially summate to induce an action potential (spike or firing) (Palmer et al., 2014). Since postsynaptic potentials from remotely located spines attenuate as they arrive at the soma, more inputs will be required to fire a neuron. Assuming that a cortical pyramidal neuron on average has nearly 3 × 10^3^ to 3 × 10^4^ spines and that a summation of nearly 100 inputs can fire a neuron, nearly 1.04 × 10^189^ to 4.68 × 10^289^ sets of combinations of inputs can fire that neuron respectively. Due to the all or none phenomenon, the firing of a neuron is not affected by inputs contributing to either sub- or supra-threshold activation of that neuron. These make neuronal firing highly non-specific with regard to the inputs and cannot be taken as specific learning-induced signature-changes. This is another constraint to arrive the correct solution. This directs the search towards a mechanism at the presynaptic terminals at the ends of converging axons and naturally towards their respective synapses.

**Figure 1.**
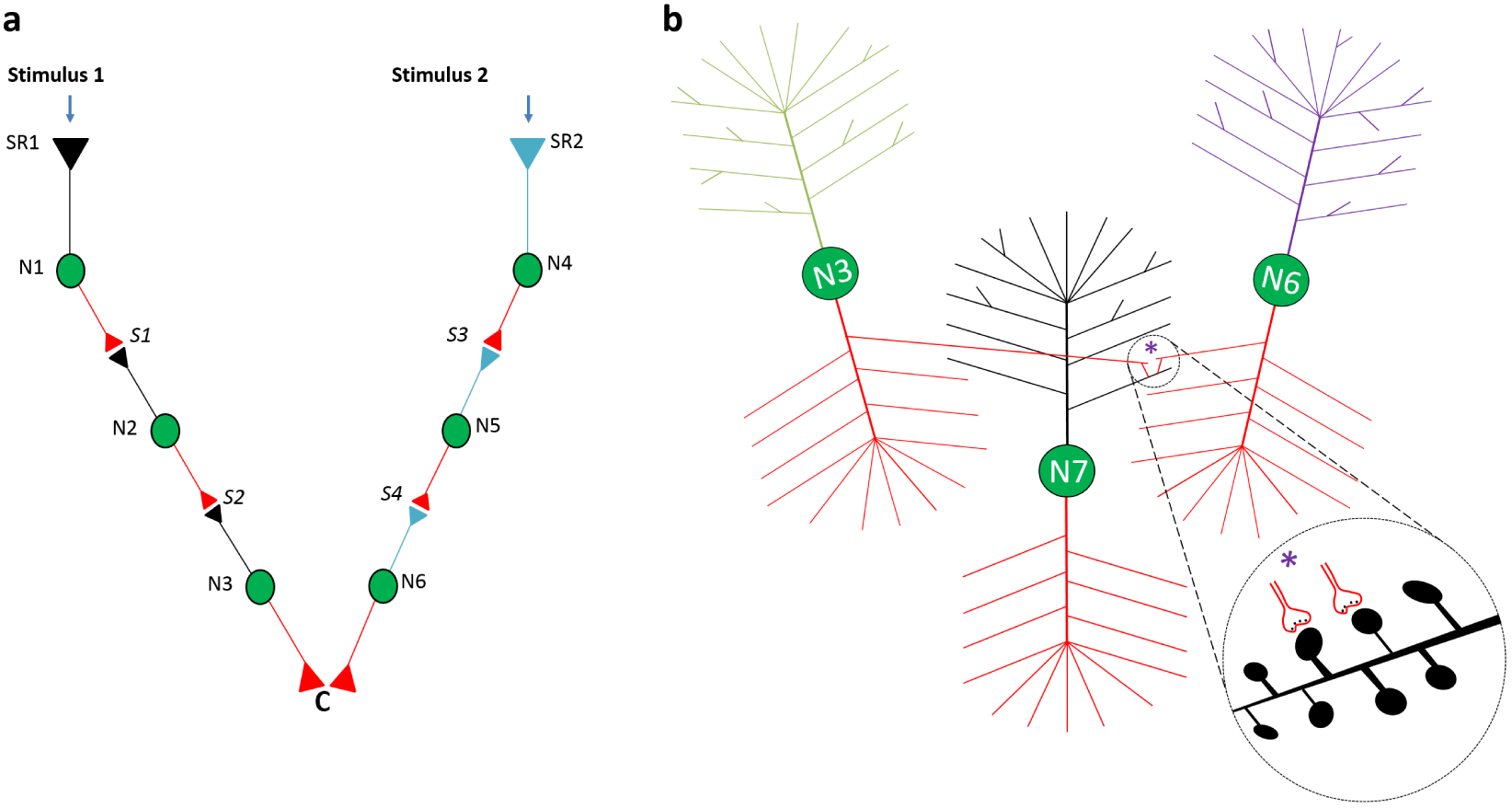
Location of convergence of sensory inputs is ideal for learning-induced change. a) Two sensory stimuli 1 and 2 arriving at sensory receptors SR1 and SR2 respectively propagate through three neurons (N1, N2, N3 and N4, N5, N6) and two synapses (S1, S2, and S3, S4) each to arrive at the location of convergence C. Changes at the synapses along their pathways (S1, S2, and S3, S4) do not provide a mechanistic explanation of how one stimulus can evoke the internal sensation of memories of the second stimulus following associative learning. Changes at the location of convergence of two stimuli (marked C) are expected to produce learning-induced changes. b) Can the converging inputs from neurons N3 and N6 synapse on to the neighboring spines of neuron N7 (magnified view in inset) for making the learning-induced change? Since the mean inter-spine distance is even more than the mean spine diameter, either a direct physical interaction or a mechanism through the ECM volume are not feasible. A mechanism through the dendritic shaft providing specificity between the spines in an electrically-isolated manner is also not feasible. The single output neuron (N7) will stop providing specific outputs associated with each of the learned stimuli. Therefore, this is not a solution.

Between what locations of the synapses of the converging inputs do learning-induced changes occur from which the cue stimulus can induce internal sensation of memory? Since neurotransmission at chemical synapses occurs in one direction, the activation of the dendritic spine can be viewed equivalent to the activation of the synapse. This indicates that if the cue stimulus can activate the postsynaptic terminal (spine) of the second stimulus, it is likely to induce a unit of the internal sensation of memory. How can the spines of the converging inputs interact? Can the converging inputs synapse on to the adjacent spines on one neuron (Fig.1b)? Dendritic shafts lack electrically-isolated conducting cables between the spines, making this unsuitable. Furthermore, since the mean inter-spine distance is even larger than the mean spine diameter (Konur et al., 2013), a mechanism through the extracellular matrix (ECM) is not possible. Moreover, it will not be possible to maintain the specific outputs associated with each of the associatively-learned sensory inputs after the level of this neuronal order, which is essential for producing motor outputs of the second stimulus by the first (cue) stimulus and vice versa. Due to the above reasons, the converging inputs from the associatively-learned sensory stimuli are expected to synapse on to the spines that belong to different neurons as a rule (Fig.2a) (Vadakkan 2016a). There could be exceptions.

**Figure 2.**
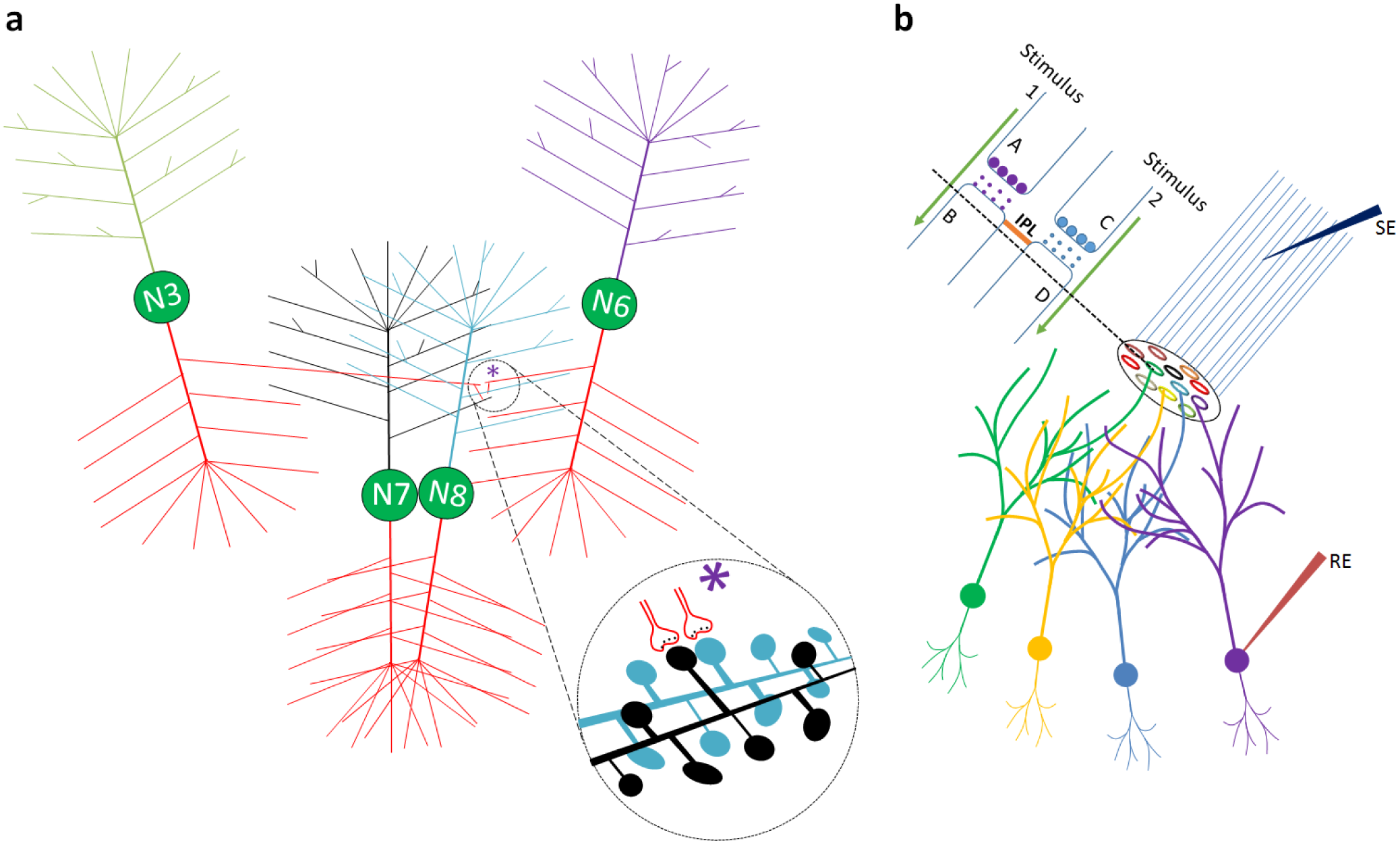
Correct level of interaction between the spines of different neurons. a) Associative learning stimuli arrive through neurons N3 and N4. Learning-induced changes are expected to take place through an interaction between the spine heads of the spines that belong to two different neurons N7 (in black) and N8 (in blue) (magnified view in the inset). This can preserve specificity of the input stimuli even after the level of their interaction by retaining the specific responses at the higher neuronal orders. b) Cross-section through an islet of inter-LINKed spines (in a circle) that belong to different neurons. One IPL is shown between the spines B and D. IPL can be formed either during learning between the readily LINKable spines or by artificial stimulation using a stimulating electrode (SE). In physiological conditions, a stimulus arriving at the islets can induce related semblances and also provide common response-motor action. Islets that are continuously activated secondary to background stimuli and shared physical properties of the items and events from the environment and can contribute to C-semblance. In contrast, large islets that are artificially-induced during LTP induction can provide a route for the regular stimulus to traverse through and reach at the recording electrode showing LTP. IPL: Inter-postsynaptic functional LINK.

The dendritic arbors of the neurons of each neuronal order overlap, including those at the locations of convergence of sensory inputs. This leads to the interaction between the abutted spine heads of different neurons during learning by the formation of specific inter-postsynaptic functional LINKs (IPLs) (Fig.2b). The term “functional” indicates that it forms as a function of the simultaneous activation of two spines at the time of associative learning and is reactivated as a function of activation of either one of the spines by the corresponding cue stimulus. The miniature potentials generated by the quantal release depolarize local areas of the spine head. Postsynaptic potentials that propagate towards the dendritic branch undergo resistance at the spine neck (Koch and Poggio, 1983; Wilson, 1984). Due to the above two reasons, the spine head region where the depolarization occurs maximally can be taken as a full-fledged, uniform, reliable location of significance where IPL can take place. The extent of the coverage of astrocytic processes over the perisynaptic area varies widely (Bernardinelli et al., 2014). Of the 57 +/-11% of synapses in the stratum radiatum of the hippocampal CA1 area that are covered by astrocytic processes, the coverage extents only less than half of the perisynaptic area (Ventura and Harris, 1999). This allows formation of the IPLs through the rest of the postsynaptic area. Continued learning leads to inter-LINKing of additional spines to the existing inter-LINKed spines, which leads to the formation of “islet” of inter-LINKed spines (called as islet) (Fig.2b). As learning continues, the size of the islet increases.

A cue stimulus is expected to reactivate the IPLs and induce bits of virtual internal sensory units that can be explained in terms of cellular hallucinations (Minsky, 1980) reminiscent of the arrival of the real stimulus from the learned item or event. What specific features permit reactivation of an IPL to induce cellular hallucinations? In the background state, continuous quantal release of neurotransmitter molecules continuously depolarizes the spine-head locally (and is thought to contribute to the miniature postsynaptic potentials). Occasionally, the arrival of an action potential at the presynaptic terminal leads to the release of a volley of neurotransmitter molecules that depolarizes the spine strongly, resulting in postsynaptic potential, which propagates towards the soma. By default, the activation of the spine-head always occurs from the presynaptic terminal. In this context, any “incidental” lateral activation of the spine-head is expected to induce the virtual internal sensation (semblance) of the arrival of activity from its presynaptic terminal as a systems property (Fig.3) (Vadakkan 2007; 2013). The incidental nature of the lateral activation can be maintained by limiting their number of occurrences by enforcing a state of sleep (Vadakkan 2016c). The sensory meaning of a unit of internal sensation can be determined by searching for the hypothetical packets of minimum sensory inputs from the environment that can normally activate the inter-LINKed spine, which are called semblions. Since this requires a search from the inter-LINKed spine towards the direction of the external environment in a retrograde direction, this constitutes the examination from a first-person frame of reference and was explained previously (Vadakkan, 2013) (Fig.3). The natural computational product of all the semblions forms memory. The induction of semblions occurs as a systems property where the lateral activation through the IPL contributes to one of the vector components of the extracellularly-recorded oscillating potentials of a specific frequency.

**Figure 3.**
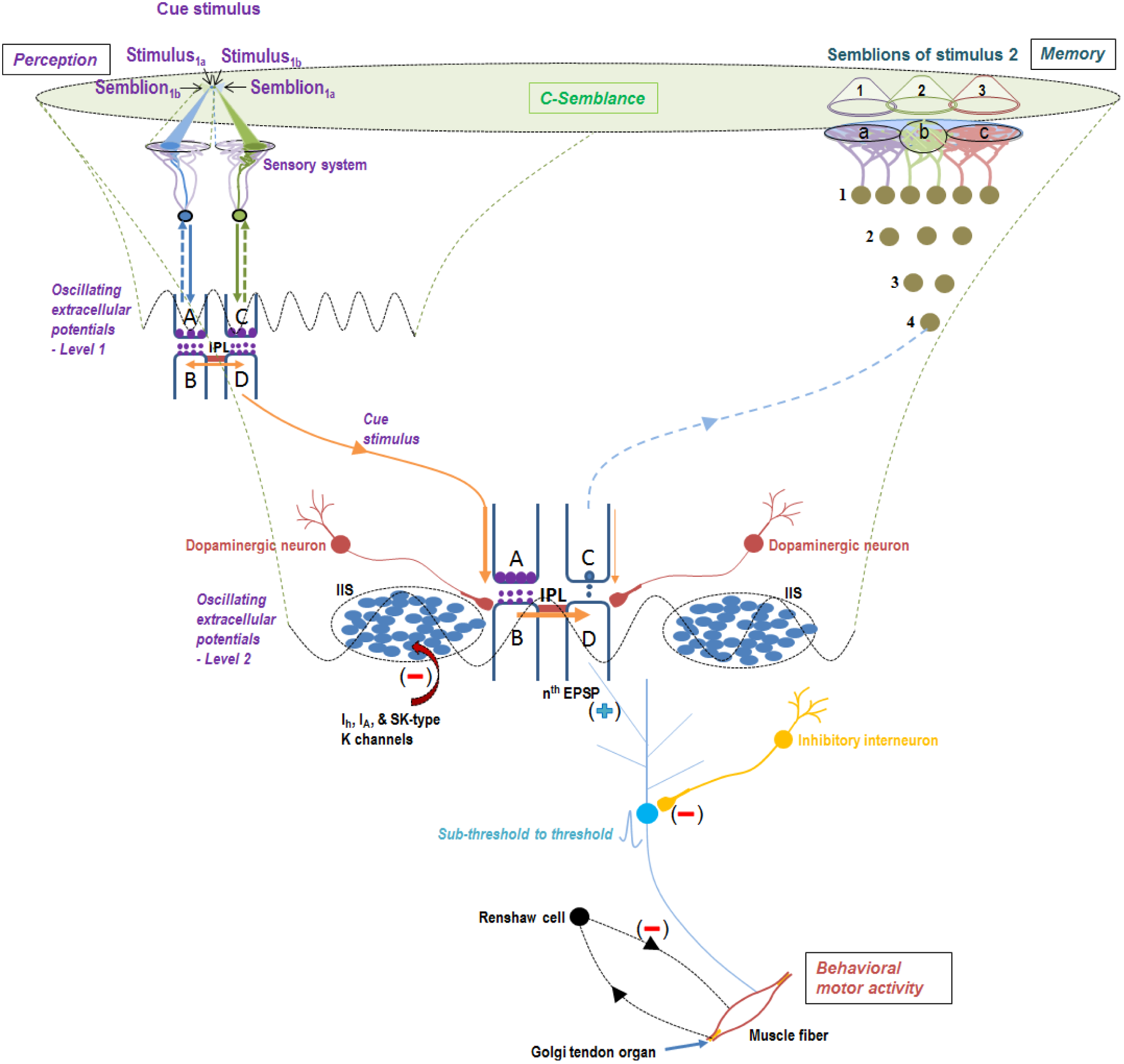
A framework for the functional organization of the nervous system using the basic structure-function unit and its modifications. On the left upper corner, the cue stimulus induces its percept at the corresponding sensory cortex. Further propagation of activity to the location of its previous convergence with a second associatively learned item leads to the activation of synapse A-B (middle area of the diagram) and an incidental reactivation of the previous associative learning-induced IPL B-D. This elicits the internal sensation (semblance) of memory of a second associatively-learned stimulus as a systems property, which is shown at the right upper corner. The sensory identity of the semblance for memory is estimated by identifying the minimum sensory stimuli that can activate presynaptic terminal C. This is derived by extrapolating from the presynaptic terminal C through the synaptic and IPL connections towards the sensory receptor level to identify the sets of minimum numbers of specific sensory receptors and the corresponding sensory stimuli (semblions). The perpendicular direction of the IPL reactivation compared to that of the synaptic transmission contributes to a vector component of the oscillating extracellular potentials. Any abnormal excitability of terminal dendrites is prevented by the activation of I*_A_*, I*_h_*, and conductance through SK-type potassium channels (Tang and Thompson, 2012). Potentials arriving at inter-LINKed spine D propagate to the soma of neuron (N) of postsynaptic terminal D. If neuron N or one of its higher order neurons is a motor neuron, then it can lead to a matching behavioural motor action. The synaptic transmission and activation of islets of inter-LINKed spines contribute to the vector components for the oscillating extracellular potentials that will activate several upstream neurons to remain at subthreshold levels. When the arrival of a cue stimulus provides additional potentials to activate a neuron held at sub-threshold state, it will lead to an associated behavioral motor action. These motor neurons are fine-regulated by inhibitory interneurons to further control its outputs. The motor response is perceived by the system in the form of proprioception or hearing that provides feedback about the motor action taken in response to the perceived cue stimulus. The net semblances from shared physical properties provide a framework for the background state of C-semblance for consciousness. Circles: in army green - excitatory neurons; in brick red - dopaminergic neurons; in yellow - inhibitory neurons. EPSP: Excitatory postsynaptic potential. IIS: Islet of inter-LINKed spines.

The nature of different types of IPLs was explained previously (Vadakkan, 2016a). Water of hydration in the inter-postsynaptic ECM volume prevents any electrical contact between the membranes. Since high energy is required to overcome the water of hydration (Leikin et al., 1987; Cohen and Melikyan, 2004; Chao et al., 2014). this type of IPL reverses back quickly and can provide a suitable mechanism for working memory. Motivation is known to release dopamine at the dopaminergic nerve terminals to the excitatory synapses, which is known to cause enlargement of the spines (Yagishita et al., 2014). If these enlarging spines are the abutted spines where the converging associatively-learned stimuli arrive, then it can result in the persistence of the IPLs for a long-period of time, increasing the probability for the latters stabilization. Since the GluR1 AMPA receptor subunits are located at the spine head locations 25nm from the synaptic margin (Burette et al., 2012), they are the likely locations where those subunit-containing vesicles get externalized resulting in reorganization of the spine membranes. This can favor IPL formation. Further enlargement of the lateral aspects of the spines can lead to hemifusion of the abutted spines, stabilization of which can explain long-term memory. Slight modifications of the formation of units of internal sensations can provide a framework for a mechanism of perception (Vadakkan, 2015b) and other higher brain functions (Fig.4).

**Figure 4.**
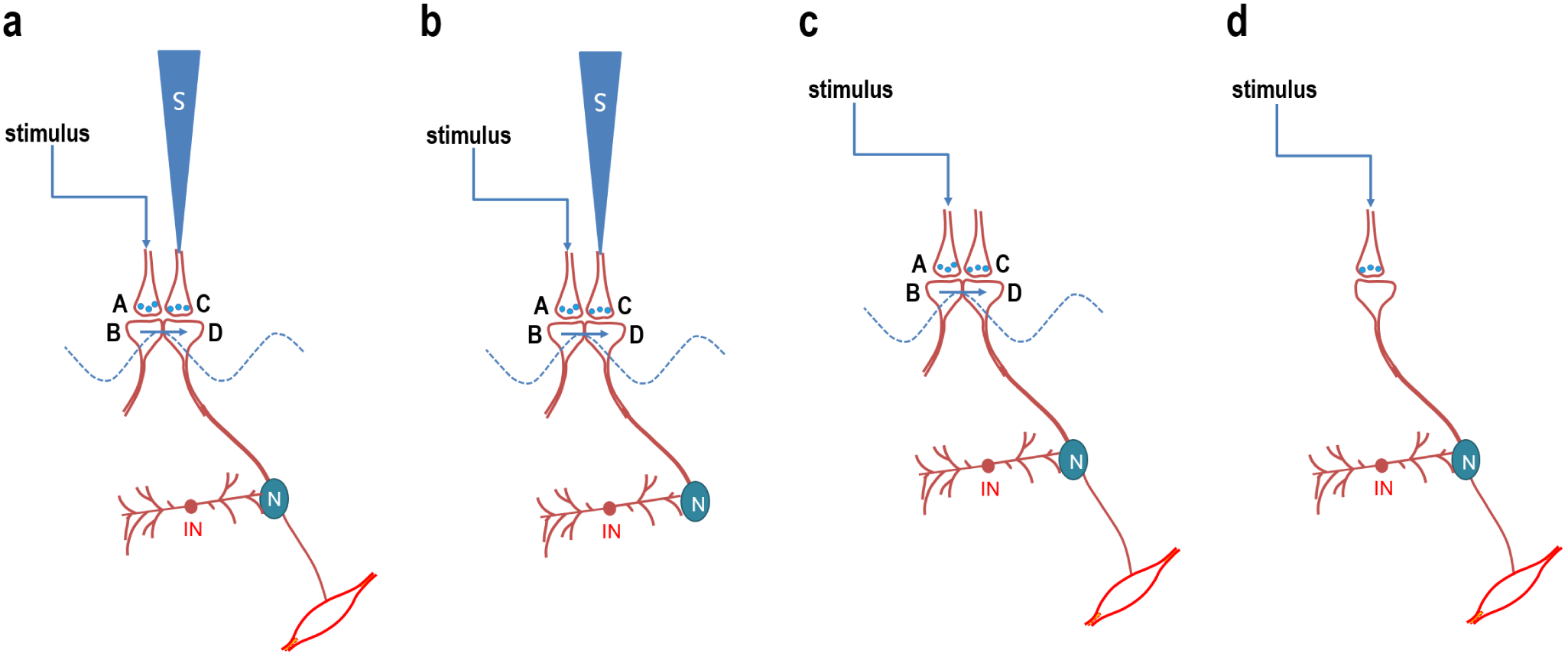
Different types of functional units of the nervous system. a) A complete unit action with the capability to induce units of internal sensation and the potential through inter-LINKed spine D can activate a motor neuron by overcoming the inhibitory actions. b) A partial unit with the capability to induce units of internal sensation and the potentials arriving through inter-LINKed spine D are inhibited by inhibitory inputs preventing motor actions either by innate mechanisms or are acquired through additional learning. A variant of these units are units that induce only internal sensations but do not carry the discriminatory activities of different stimuli forward. Dendritic excrescences are examples. c) A partial unit action that takes place in the presence of oscillating extracellular potentials but without the capability to induce units of internal sensation. The potential through inter-LINKed spine D overcomes the inhibitory inputs to activate a motor neuron. These types of units can be found in the spinal cord. d) A direct synaptic activity that leads to the activation of a motor neuron by overcoming the inhibitory inputs. N: Excitatory or inhibitory neuron. IN: Inhibitory neuron. A and C: Presynaptic terminals. B and D: Postsynaptic terminals (spines).

### 2.3 All-inclusive circuitry

IPL mechanism can explain a large number of diverse functions at various levels. These can take place within the framework of a functional organization consisting of different modifications of the basic structure-function units. a) IPL mechanism operates in unison with the synaptically-connected neuronal circuitry. b) At the glutamatergic synapses, in the absence of dopaminergic inputs, IPL formation and reversal may depend on the AMPA receptor vesicle exocytosis and endocytosis producing reorganization of the lateral aspects of the spines at physiological time-scales. c) All the biochemical changes occurring slower that the physiological time-scales can be viewed as cellular homeostatic mechanisms preparing the synapses for future operations. d) Potentiation of both synaptic transmission and AMPA receptor-mediated excitatory postsynaptic potentials (EPSPs) by muscarinic M1 acetyl choline receptors (Dennis et al., 2016) can be explained in terms of the universal IPL-mediated mechanism. e) The circuit mechanism can introduce control of motor actions through additional associative-learning events or introduce a delay in motor response. f) The IPL-mediated mechanism can explain how regenerative dendritic events take place and their relationship with place cell firing properties (Sheffield and Dombeck 2015). g) If a specific cue stimulus x was associated with two different sensory stimuli y and z in the past, the presence of x can lead to the induction of semblances for both y and z. Induction of these two semblances simultaneously can generate a hypothesis that y and z are inter-related. If two items get associated through a series of steps, but short of one step in between, the system may get the internal sensation (imagination) about this missing step. h) A lack of sensory inputs during sleep will lead to a reduction in spine head size, which can increase the glymphatic flow (Xie et al., 2013). i) Primitive motor actions necessary for survival can be formed by an innate mechanism of formation of IPLs, that are evolutionarily acquired. j) Insertion of new neuron at one neuronal order of the circuitry with repetition of learning or related learning can lead to the formation of increased number of IPLs at the level of higher neuronal order. This can explain augmentation of learning and consolidation of memory (Vadakkan, 2011a). k) In the absence of repetition of learning or related learning or cue-induced memory retrieval after the initial learning event, insertion of new neurons at one neuronal order of the circuitry will dilute the net specific semblance in response to a cue stimulus. l) The correlation between memory and LTP can be explained in terms of the formation islets of inter-LINKed postsynaptic terminals (Vadakkan, 2013; Vadakkan, 2016e). m) Kindling is induced by stronger stimulation conditions than LTP and shows several similarities with human seizure disorders (Bertram, 2007; Vadakkan, 2016b) The expected non-reversible inter-spine fusion events can support the finding that spatial memory performance gets disrupted by kindling and not by hippocampal CA1 LTP (Leung and Shen, 2006). n) The finding that perineural net proteins in the ECM space at the CA2 region blocks LTP (Carstens et al., 2016) and CA2 region is uniquely resistant to seizures (Hatanpaa et al., 2014) provide support for the IPL mechanism. o) Changes at the level of the spines (Fuhrmann et al., 2007) that eventually lead to IPM fusion and spine loss (Herms and Dorostkar, 2016) can explain different neurodegenerative disorders. Over-activation of the NMDA receptors (Zeron et al., 2002) and excessive dopamine in the early stages of Huntington‘s disease (Cepeda et al., 2014) are some of the causes. Neurodegeneration can lead to both cognitive and motor defects (Vadakkan, 2016d). p) Formation of pathological mis-LINKs between spines can lead to hallucinations (Vadakkan 2012a). q) Since many physical properties of the items in the environment are shared, the stimuli arriving from them and from within the body are expected to continue to activate a large number of shared islets of inter-LINKed spines. The lateral spread of activity through these IPLs contribute to the normal frequency of the oscillating extracellular potentials and the induced semblances are expected to contribute to the background C-semblance (Vadakkan, 2010). The C-semblance forms a background matrix or medium upon which specific IPLs are formed during learning, which involves the binding of different sensations in the nervous system (von der Malsburg, 1999). Maintaining this background state is also required for a specific cue stimulus to induce specific semblances for memory. It also maintains several neurons at subthreshold-activation levels, which can be activated using minimal potentials arriving through the reactivation of IPLs for executing behavioral motor actions. A dendritic spike, which is a synchronous activation of 10 to 50 neighboring glutamatergic synapses triggering a local regenerative potential (Antic et al., 2010) and occurring *in vivo* (Cichon and Gan, 2015) may contribute to the C-semblance. Since consciousness is considered the most important binding process (Roskies, 1999), the intrinsic nature of C-semblance can provide required features of a framework for consciousness (Vadakkan, 2010; Fekete et al., 2016). r) Unconscious motor responses occur in response to common stimuli to which the system gets exposed routinely. This is because the common stimuli reactivate IPLs within the islets of inter-LINKed spines such that the induced-semblances that are integrated with the C-semblance do not form a specific semblance. This can lead to the unconscsious occurrence of common stimuli-induced motor responses. s) The formation of a large number of non-specific IPLs triggered by membrane alterations by lipophic anesthetic molecules can explain the effect of anesthetic agents (Vadakkan, 2015a).

A comparative circuitry of the synaptically-connected neural circuitry in *Drosophila* can accommodate all the IPL-mediated mechanisms (Fig.5). The synaptically-connected neural circuitry of the fly can accommodate features necessary for the first-person internal sensation of olfactory perception in a background state of internal sensation of awareness (Vadakkan 2015b). The systems requirement for sleep (Vadakkan 2016c) for generating various internal sensations can be observed in the fly as sleep-like states (Hendricks et al., 2000). The higher neuronal orders from the olfactory circuitry that converge with that of other sensations such as vision can form IPLs during associative learning. In summary, the axonal terminals of similar types of olfactory receptor neurons (ORNs) synapse to the dendritic spines of the sister projection neurons (PNs) within a specific glomerulus in the antennal lobe. Several IPLs can be formed between the spines of PN neurons within a glomerulus. During the baseline awake state, the ORN neurons having a baseline firing frequency of 8 spikes/s activate several IPLs within the glomeruli that can induce C-semblance for awareness. When the fly is exposed to a unique odorant, it can stimulate specific types of ORNs and activate the spines of the PNs within a specific glomerulus. This leads to reactivation of the IPLs from both sides, which will lead to the generation of stimulus-semblion U-loops to form the perceptons. The net semblance of all the perceptons can form the percept of a specific smell (Vadakkan 2015b). While one glomerulus is activated, inhibitory local interneurons inhibit all the remaining glomeruli (Hong and Wilson, 2015), enabling specificity of the percept for a particular smell. The odor-induced ORN-PN synaptic activation and the spread of potentials through the IPLs within the glomerulus contribute to the vector components of the oscillating potentials, which will lead to oscillatory synchronization of spiking in groups of PNs found to be associated with fine-odor perception (Stopfer et al., 1997; Laurent, 1999). PN axonal terminals bifurcate to innervate two distinct regions - the mushroom body (MB) and lateral horn (LH). In the MB, they synapse to Kenyan cell (KC) neurons. The output neurons from the MB are the mushroom body output neurons (MBONs). Following the perception of smell in a conscious state, the stimuli can reach the locations of convergence with other sensory stimuli, such as the sight of food, which can lead to changes necessary for associative learning. The axonal terminals from neurons of different sensory inputs such as vision and mechano-sensation synapse to the spines of the MBONs. Therefore, simultaneous activation of olfactory and other sensory stimuli can lead to reversible, yet stabilizable IPLs between the spines of the MBONs during associative learning. At a later time when one sensory stimulus arrives, it can induce the internal sensation of memory of the associatively-learned second stimulus.

**Figure 5.**
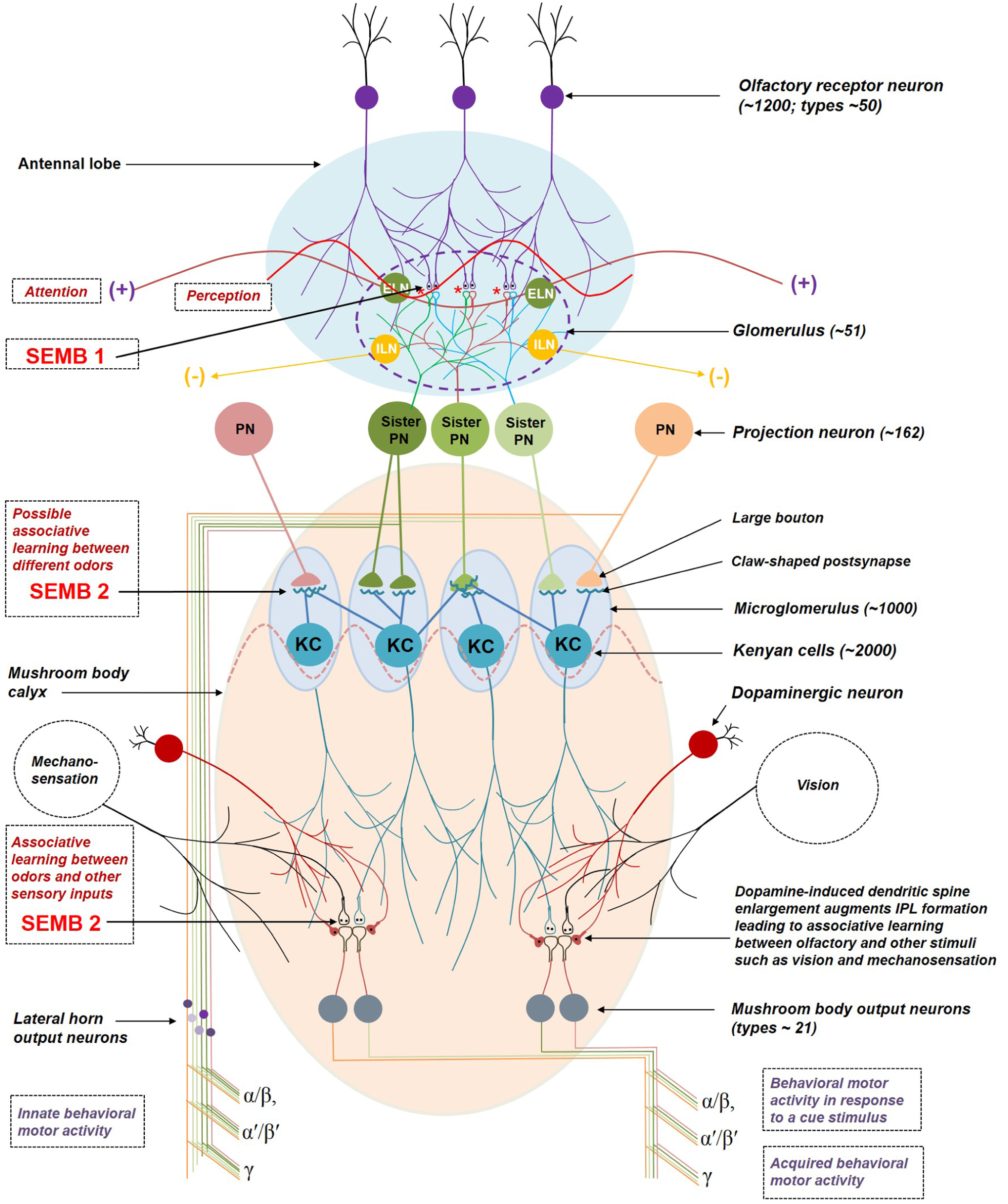
The synaptically-connected nervous system of *Drosophila* olfactory nervous system provides both first-person sensible and third-person observable functions. The number of neurons or processes is given in the brackets. The binding of an odorant to the receptors on a set of ORNs results in the activation of a specific set of PNs whose dendritic arbors are confined to specific glomeruli. All of the ORNs that express a given odorant receptor converge onto the same glomerulus in the antennal lobe. The presynaptic terminals of ORNs synapse with the dendritic spines of 3 sister PNs within the glomerulus. In the absence of ligands, ORNs make 8 spikes per second continuously. Nearly 3 synchronous unitary ORN synaptic inputs activate a PN to elicit one spike. At rest, the activity from ORNs arriving at one glomerulus spreads to other glomeruli through excitatory cholinergic local interneurons. The arrival of one type of ligand (odorant molecule) activates one type of ORNs, which inhibits all other glomeruli by activation of inhibitory local interneurons. The PN projects axons that bifurcate to innervate two distinct brain regions, the MB and the LH. PN axons innervate the MB by terminating in large boutons and synapse with one claw each of many Kenyan cells (KCs). A single bouton connects to multiple KC claws; but each claw synapses with only one PN bouton. PNs from different glomeruli synapse to the claws of the KCs by chance alone. Odorant-evoked local field potential oscillations are seen in the MB. Each KC projects an axon to one of the three different classes of MB lobes, *α/β, α′/β′, or γ*, where it synapses upon a relatively small number of MBONs. The general architecture of the fly nervous system is comparable to that of the mammals as follows. 1) PNs are interconnected through the ELNs and the basal activation spreading through them is observed as oscillatory local field potentials. Non-specific background semblances induced by re-activation of the IPLs between the spines of the PNs in glomeruli and KC neurons in the microglomeruli induce a net non-specific background semblance, which is expected to give rise to the C-semblance responsible for attention/consciousness. 2) Stimulus-semblion U-loops formed at the IPLs of the initial sensory neuronal orders can provide the formation of perceptons to perceive olfactory sensory stimuli. 3) IPLs are expected to form during associative learning at locations of convergence of sensory stimuli that can be re-activated to form semblances for memory. IPLs can be formed between the spines of the MBONs during associative learning between an olfactory stimulus and other sensory stimuli like vision or mechano-sensory stimuli. The dopaminergic neuronal terminals synapsing on to the spines of the MBONs can promote the latters enlargement and facilitate the formation of IPLs. This can augment associative learning. The arrival of an olfactory stimulus can then lead to the reactivation of these IPLs, leading to the internal sensation of memory of associatively-learned sensory stimuli and vice versa. 4) The oscillatory neuronal activity of the KC neurons is expected to keep many of the MBONs at sub-threshold level of activation. Olfactory stimuli, along with inducing semblances for the associatively-learned stimuli, can provide additional potentials to the specific MBONs for their firing leading to behavioral motor action. It is expected that inhibitory interneurons are involved in fine-regulating the behavioral motor action at the level of the MBONs. 5) PN connections to the lateral horn output neurons (LHONs) are fixed and are thought to mediate innate behaviors through LHONs. It can be expected that the IPLs between the spines of PNs within the glomerulus are stabilized through innate genetic mechanisms. Reactivation of IPLs induce a percept concurrent with triggering of motor activity of flight. SEMB 1: semblance at the spines of PN; SEMB 2: semblance at the spines of KCs; SEMB 3: semblance at the spines of MBONs.

PN axons terminate in large boutons (Marin et al., 2002; Wong et al., 2002), each of which synapses to multiple KC claws (Yasuyama et al., 2002). Each KC neuron has an axonal projection to one of the three different classes of MB lobes *α/β, α′/β′, or γ* where it synapses upon a relatively small number of MBONs that are anatomically segregated and are responsible for the different forms of learned behavior (Tanaka et al., 2008; Sjourn et al., 2011). Nearly 2,000 KCs converge onto a population of only 34 MBONs that are of 21 distinct cell types (Aso et al., 2014). Since some of the MBONs receive connections from almost all the KCs (Aso et al., 2014), it indicates the presence of stereotyped behavioral activity in response to most of the odors. Dopamine is associated with motivation-promoted associative learning (Wise, 2004). In *Drosophila*, the role of dopamine in appetitive and aversive memories is well-documented (Waddell, 2010; Liu et al., 2012). Dopamine neurons in the protocerebral anterior-medial cluster have their axonal terminals in the MB. The presynaptic boutons at the axonal terminals of these dopaminergic neurons synapsing on the spines of the MBONs can lead to the enlargement of the latter, which can promote IPLs between them. Thus, motivation-promoted associative learning can possibly occur through the formation of IPLs between the spines of the MBONs through dopamine-induced spine enlargement (Yagishita et al., 2014). CO_2_ is a repellant and it specifically activates one specific glomerulus (Suh et al., 2004), which is expected to induce semblance for an aversive internal sensation.

Odorant-evoked local field potential (LFP) oscillation seen in the MB (Paulk et al., 2013) likely results from the lateral spread of activity through the IPLs between the claws of the KC neurons. This can form one vector component of oscillating potentials and contribute to the phase-locking with the LFP oscillations in the MB (Tanaka et al., 2009). LFP oscillations may also lead to the induction of a non-specific set of semblances at the KC claws that contribute to the C-semblance for fly attention. The baseline oscillating potentials can lead to sub-threshold activation of several upstream MBONs that can allow these neurons to fire at the arrival of one or a few potentials through an IPL concurrent with the internal sensation of memory of smell evoked by a visual stimulus and vice versa. Since light travels faster than smell, this allows motor action at the slightest visual sensory stimulus and can provide survival advantage. Similarly, smell arriving from hidden locations can offer information about the items in the environment. The motor action of flight will provide proprioceptive feedback stimuli to the nervous system to optimize its functional efficiency. The stereotyped combinations of outputs from glomeruli that are co-activated by the odors are thought to be responsible for the innate behaviours (Fisek and Wilson, 2014). The invariant nature of the LH is thought to mediate these innate behaviors.

### 2.4 Multiple inter-connected triangulations provide proof

Triangulation is a powerful technique for validating the interrelationships within a system through cross verifications. As the triangulated findings from one level can be further triangulated with the findings from different levels, it increases the strength of the proof for the first-principle from which all the findings arise (Supplementary Information IV). Triangulations between the features of “loss of function” states of these functions further strengthen the evidence.

### 2.5 Functional efficiency of the system

The operations of the system are taking place at the level of the IPLs that are located between the layers of neuronal soma. What are the limiting factors for the efficient operation of the system? These include a) factors that guide the systems property of the induction of internal sensations such as frequency of oscillating extracellular potentials, b) number of non-LINKed abutted spines at the locations where associatively learned sensory stimuli converge, and c) the ability to keep the neurons of these spines just below the threshold for spiking during the background state of oscillating extracellular potentials occurring at a particular frequency. The contribution of the overlapping semblances to the sensitivity and specificity of the memory will vary at different orders of IPLs.

## 3 Discussion

The simple nature of the unique mechanism that can explain and interconnect findings from large number of levels shows its feasibility to be present throughout the animal species. The technically-challenging imaging of the real-time occurrence of different types of IPLs that span areas nearly 10nm2 can provide proof. IPL formation and function do not require the presence of any distinct cortical areas, supporting the previous observations (Gntrkn and Bugnyar, 2016). The functional attributes of the IPL-operated functional units can accommodate heterogeneity of neuronal types within the nervous system. Pyramidal neurons of human cortical layer L2 and L3 have 3-fold increase in dendritic length and branch-complexity compared with those of macaque and mouse (Mohan et al., 2015). This may explain the presence of more abutted spines that can be inter-LINKed and also form large islets. Facilitation of IPL formation by dopamine and control of the outputs through inhibitory neuronal activity (Palmer et al., 2012) provide regulatory features. Once humans learned to protect themselves by the use of tools, improved inhibitory features to control reflex actions were likely added to the circuitry, which eventually led to the refinement of behavior during the induction of various internal sensations in response to cue stimuli. Any non-physiological stimulation of the neurons can lead to the formation of non-specific IPLs at higher neuronal orders, inducing non-specific internal sensations and motoric effects.

The IPL-mediated mechanism is expected to allow the system to operate autonomously and perform unsupervised learning and reasoning. Feasibility of the synaptically-connected circuitry in the flys nervous system to accommodate IPLs to explain the required functional features at different levels indicates the universal presence of the IPL mechanism across the animal species. The property of specific neuronal circuit organization that is expected to induce first-person internal sensations of various higher brain functions matches with the expectations of an ideal system through the reactivation of K-lines that can produce biological memory (Minsky, 1980). The IPL-mediated mechanism can be replicated in engineered systems (Vadakkan, 2011b), which can be used to test and study the nature of internal sensations of various higher brain functions (Vadakkan, 2012b). The gold standard test of this theoretically-feasible mechanism requires replication in engineered systems, which simultaneously provides an opportunity to develop humanoid artificial intelligence. The algorithms of the natural computations of the units of internal sensations for a given system is expected to be discovered while undertaking this replication.

## Acknowledgements

This work was supported by Neurosearch Center, Toronto. I would like to thank Selena Beckman-Harned for reading the manuscript.

## Conflict of interest

U.S. patent: number 9,477,924 pertains to an electronic circuit model of the inter-postsynaptic functional LINK.

## Supplementary Information

### I. List of features requiring an explanation from a mechanism of nervous system function

1. Mechanism for the formation of virtual first-person internal sensations 2. Induction of internal sensations at physiological time-scales 3. Instant access to very large memory stores (Abbott, 2008) 4. Induction of internal sensation of specific memory in response to specific cue stimulus 5. Reversible and stabilizable mechanism for storing memories 6. Absence of cellular changes during memory retrieval 7. Instant access to very large number of memory stores 8. Working, short- and long-term memories are induced from the same mechanism; but retained for different lengths of time following the associative learning 9. Half-life of the stabilization mechanism determining the duration of memory 10. Operates in unison with the known synaptically-mediated circuit operations 11. Storage of very large number of memories 12. Motivation-promoted learning explainable by a mechanism by the released dopamine 13. Formation of a specific memory in response to a specific cue stimulus 14. Retrieval of memories after very long period of time following the learning 15. Ability to store new memories without needing to overwrite the old ones 16. Feasibility to forget previously-learned items or events 17. Consolidation of memory 18. Reconsolidation occurring by exposing to the cue stimulus after a period of time 19. Mechanism to use schemas inter-changeably (Tse et al., 2007) 20. Mechanism providing source for potentials contributing to the vector components of the oscillating extracellularly-recorded potentials 21. Relationship of different higher brain functions with the frequency of extracellularly-recorded oscillating potentials 22. Ability to make hypothesis (Abbott, 2008) 23. Correlation between LTP and surrogate markers of retrieved memory 24. Relationship between LTP, kindling and seizures 25. Reduction of LTP by the blockers of postsynaptic membrane fusion (Lledo et al., 1998) 26. Induction of LTP at the CA2 area of the hippocampus becomes possible by the removal of the perineural net proteins chemically (Dudek et al., 2016) 27. Resistance of the CA2 area of the hippocampus for seizures (Hatanpaa et al., 2014) 28. Dementia in Alzheimers disease 29. Cognitive defects in several viral infections, especially by herpes simplex virus (HSV) 30. Seizures in HSV viral encephalitis (Michael and Solomon, 2012) 31. Place cell firing in response to specific spatial stimulus 32. Firing of motor neurons that lead to behavior indicative of retrieval of a specific memory 33. Mechanism for innate behavior 34. Framework for consciousness 35. Loss of consciousness by anesthetics 36. Effect of dopamine in augmenting anesthetic action (Segal et al., 1990) 37. Reversal of the loss of consciousness after a generalized seizure 38. Interconnection between seizures and loss of consciousness 39. Paroxysmal depolarization shift (PDS) during seizures (Johnston and Brown, 1981) 40. Internal sensation of hallucinations 41. Mechanism for hallucinations and cognitive defects in seizures 42. Loss of dendritic spines after kindling (Singh et al., 2013) 43. Heterogeneity of observations in neurodegenerative disorders (Sahin et al., 2015) 44. Indispensable need for sleep 45. Memory storage without needing to store information in any form of energy 46. Perception as a first-person internal sensation 47. Flash lag delay, apparent location of the percept different from the actual location, homogeneity in the percept for stimuli above the flicker fusion frequency and a mechanism for perceiving object borders 48. Mechanism for neurodegenerative disorders 49. Presence of comparative circuity in a remote species

## II. Comparison of solving a system of linear algebraic equations by the invention of negative integers

Solving the nervous system can be compared to solving a set of large number of algebraic equations to find the value of an unknown factor. The case of the virtual nature of inner sensation of higher brain functions can be explained by this method. Imagine that we are living in a period before the invention of negative integers. We came across a system that operates by interconnecting three unknown variables. We obtained three linear algebraic equations from the system as follows. In order to understand the system, it is required to solve the system.

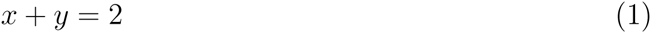

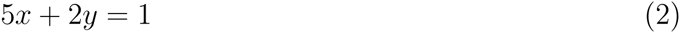

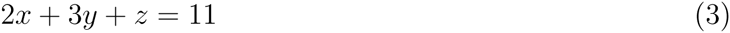

Attempts can be made to solve this system of linear algebraic equations as follows. All of the equations have the unknown factor x. The simplest two equations are (1) and (2), from which the value of x can be calculated by the method of cancellation as follows.

Multiplying (1) with 5, we get

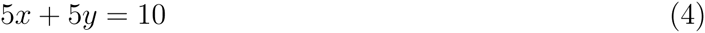

Subtracting equation (2) from equation (4), we get

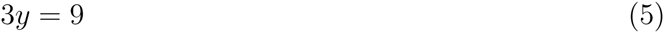

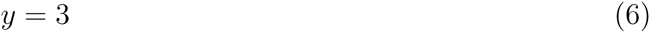

Substituting y = 3 in equation (1), we get

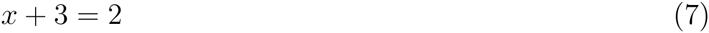

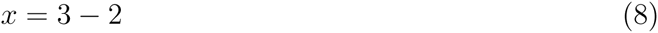

It was not possible to find the solution for x before the invention of negative integers. In other words, it was required to invent a method to subtract 3 from 2 by introducing negative integers. It was not possible to think of negative integers because human beings do not have a sensory system to sense them directly. Therefore, it is necessary to assign a virtual, physically non-existing value of -1 for x. This facilitates solving the remaining equation (3). Plotting the negative integers on the x-axis of a graph facilitated perceiving them better. Substituting the values x = -1 and y = 3 in equation (3), we get z = 4. But, we are not sure whether the value of x = -1 is correct or not. In order to verify this, it is necessary to search for more linear algebraic equations from the system. Imagine that the following equation is obtained,

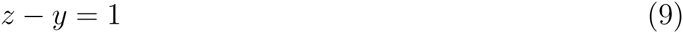

By substituting the values for z and y, it verifies the previous observations and provides some assurance that the value assigned to x = -1 is correct. By testing more equations from the system, it will be possible to confirm the value of x for the system. The virtual nature of the negative integer has similarities to observing internal sensations from a third-person view. Virtual internal sensations are essential features of all the higher brain functions. Similar to the invention of negative integers we can assign an imaginary mechanism, which can be induced from a third-person-observed cellular change. It can be verified using retrodictive approaches by using already gathered data, which will make the examination free of any experimenter bias.

## III. Solving a system requires incorporating all the variables and using at least a minimum number of functions within the system

Since a large number of findings are observed at various levels (biochemical, subcellular, cellular, inter-cellular, electrophysiological, systems, and behavior), solving the nervous system requires an approach similar to solving a system of equations in multiple variables. It is necessary to include all the variables to find the unique solution that can solve the whole system. Trying to solve the system by using a subset of equations having only a subset of variables will result in obtaining a wrong solution for the whole system. This can be explained by the following example. Let us try to solve the following set of four equations that have four variables M, P, T, and Z (Since different findings are obtained from different levels, let us assume that all the variables of the system have different values). This is similar to trying to solve the nervous system using the key features from its four levels for example, biochemical, electrical conduction properties, neuronal firing and behavior.

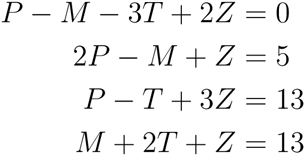

One of the solutions is M = 2; P = 1; T = 3; Z = 5 (In fact, the above set of equations has infinitely many solutions that satisfy the conditions P = (2*M* − 1)/3; *T* = (11 − *M*)/3; *Z* = (17 − *M*)/3). What does these values mean? The values of the variables M, P, T and Z strongly interconnect the set of four equations. They bind these equations together within the system. The values of the variables allow these equations to form an independent operational system if the system has only these four equations. But what happens when the system has more variables? Let us introduce few new variables K, N, O, R, S, Y by adding more equations that contain both the previous and the new variables into the system. We get a new system of equations, which is shown below. This is similar to examining additional key features from more levels of the nervous system for example, subcellular, inter-cellular, systems, long-term potentiation, extracellular potentials, and sleep. Let us assume that we have used a fair sample of features from all the different levels and this is the complete system. Let us now examine the system.

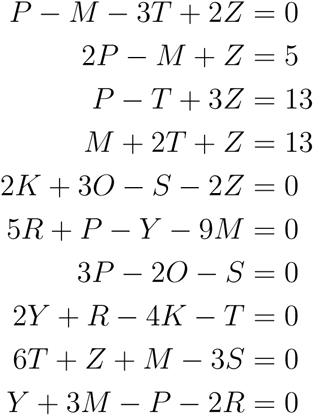

The previous solution (M = 2; P = 1; T = 3; Z = 5) for the previous system of four equations (which is a subsystem of the whole system of ten equations) will not be fitting for the whole system. Instead, their values need to change to M = 5; P = 3; T = 2; and Z = 4 (One of the solutions for other variables are K = 6; O = 1; R = 10; S = 7; Y = 8). This means that the variables have to assume new values for them to get integrated into the system to interconnect the system together. This explains that adding more variables to the system can change the solution for the system. This example highlights the fact that it is required to include all the variables by using a set of minimum number of equations to find a solution for a system. Therefore, in the case of the nervous system the solution can be arrived only by incorporating the findings from all the different levels. This is necessary to confirm that the arrived solution for the system will be able to explain and inter-connect all the findings at different levels.

## IV. Multiple inter-connected triangulations between the findings from the normal and loss of function states

**Figure 6.**
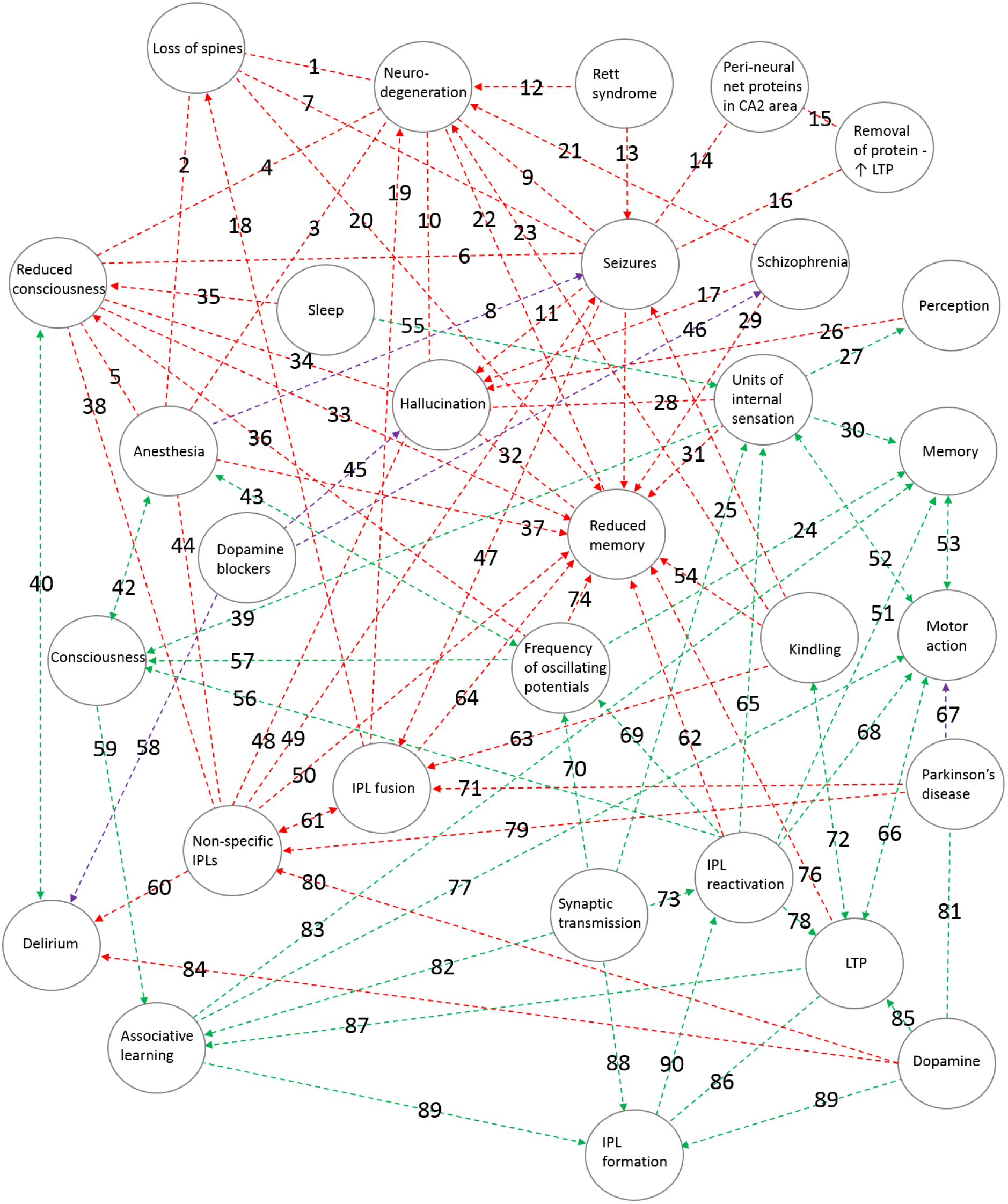
A large number of triangulations between the findings from different levels of the normal functions, loss of function states and the effect of pharmacological agents (some are left out due to overcrowding). This ability to make a large number of inter-connected triangulations supports IPL-mediated mechanisms as a first-principle of the nervous system functions. Both direct and indirect triangulations can be observed in the figure. The interconnecting lines are marked with specific numbers. Green broken lines: normal functions, Red broken lines: pathological conditions, Violet broken lines: action that reduces the effect, Single arrows: unidirectional effect, Double-headed arrow: bidirectional effect.

## References

Abbott LF (2008) Theoretical neuroscience rising. Neuron. 60:489.

Antic SD, Zhou WL, Moore AR, Short SM, Ikonomu KD (2010) The decade of the dendritic NMDA spike. Journal of Neuroscience Research. 88:2991.

Aso Y, Hattori D, Yu Y, Johnston RM, Iyer NA, Ngo TT, Dionne H, Abbott LF, Axel R, Tanimoto H, Rubin GM (2014) The neuronal architecture of the mushroom body provides a logic for associative learning. Elife. 3:e04577.

Bernardinelli Y Muller D, Nikonenko I (2014) Astrocyte-synapse structural plasticity. Neural Plasticity. 2014:232105.

Bertram E (2007) The relevance of kindling for human epilepsy. Epilepsia. 48 Suppl 2:65.

Burette AC, Lesperance T, Crum J, Martone M, Volkmann N, Ellisman MH, Weinberg RJ (2012) Electron tomographic analysis of synaptic ultrastructure. Journal of Comparative Neurology. 520:2697.

Carstens KE, Phillips ML, Pozzo-Miller L, Weinberg RJ, Dudek SM (2016) Perineuronal nets suppress plasticity of excitatory synapses on CA2 pyramidal neurons. Journal of Neuroscience. 36: 6312.

Cepeda C, Murphy KP, Parent M, Levine MS (2014) The role of dopamine in Huntington’s disease. Progress in Brain Research. 211:235.

Chao LH, Klein DE, Schmidt AG, Pea JM, Harrison SC (2014) Sequential conformational rearrangements in flavivirus membrane fusion. Elife. 3:e04389.

Cichon J, Gan WB (2015) Branch-specific dendritic Ca(2+) spikes cause persistent synaptic plasticity. Nature. 520:180.

Cohen FS, Melikyan GB (2004) The energetics of membrane fusion from binding through hemifusion pore formation and pore enlargement. Journal of Membrane Biology. 199:1.

Cragg BG (1967) The density of synapses and neurons in the motor and visual areas of the cerebral cortex. Journal of Anatomy 101:639.

Dennis SH, Pasqui F, Colvin EM, Sanger H, Mogg AJ, Felder CC, Broad LM,Fitzjohn SM, Isaac JT, Mellor JR (2016) Activation of muscarinic M1 acetylcholine receptors induces long-term potentiation in the hippocampus. Cerebral Cortex. 26:414.

Edelman S (2012) Six challenges to theoretical and philosophical psychology. Frontiers in Psychology. 3:219.

Fekete T, van Leeuwen C, Edelman S (2016) System subsystem hive: Boundary problems in computational theories of consciousness. Frontiers in Psychology. 7:1041.

Fisek M, Wilson RI (2014) Stereotyped connectivity and computations in higher-order olfactory neurons. Nature Neuroscience. 17:280.

Fuhrmann M, Mitteregger G, Kretzschmar H, Herms J (2007) Dendritic pathology in prion disease starts at the synaptic spine. Journal of Neuroscience. 27:6224.

Gntrkn O, Bugnyar T (2016) Cognition without cortex. Trends in Cognitive Science. 20:291.

Hatanpaa KJ, Raisanen JM, Herndon E, Burns DK, Foong C, Habib AA, White CL 3rd. (2014) Hippocampal sclerosis in dementia epilepsy and ischemic injury: differential vulnerability of hippocampal subfields. Journal of Neuropathology and Experimental Neurology. 73:136.

Hendricks JC, Finn SM, Panckeri KA, Chavkin J, Williams JA, Sehgal A, Pack AI (2000) Rest in Drosophila is a sleep-like state. Neuron. 25:129.

Herms J, Dorostkar MM (2016) Dendritic spine pathology in neurodegenerative diseases. Annual Review of Pathology. 11:221.

Hong EJ, Wilson RI (2015) Simultaneous encoding of odors by channels with diverse sensitivity to inhibition. Neuron. 85:573.

Koch C, Poggio T (1983) A theoretical analysis of electrical properties of spines. Proceedings of the Royal Society B: Biological Sciences. 218:455.

Konur S, Rabinowitz D, Fenstermaker VL, Yuste R (2003) Systematic regulation of spine sizes and densities in pyramidal neurons. Journal of Neurobiology. 56:95.

Laurent G (1999) A systems perspective on early olfactory coding. Science. 286:723.

Leikin SL, Kozlov MM, Chernomordik LV Markin VS, Chizmadzhev YA (1987) Membrane fusion: overcoming of the hydration barrier and local restructuring. Journal of Theoretical Biology. 129:411.

Leung LS, Shen B (2006) Hippocampal CA1 kindling but not long-term potentiation disrupts spatial memory performance. Learning and Memory. 13:18.

Liu C, Plaais PY, Yamagata N, Pfeiffer BD, Aso Y, Friedrich AB, Siwanowicz I, Rubin GM, Preat T, Tanimoto H (2012) A subset of dopamine neurons signals reward for odour memory in Drosophila. Nature. 488:512.

Marin EC, Jefferis GS, Komiyama T, Zhu H, Luo L (2002) Representation of the glomerular olfactory map in the Drosophila brain. Cell. 109:243.

Minsky M (1980) K-lines: a theory of memory. Cognitive Science. 4:117.

Mohan H, Verhoog MB, Doreswamy KK, Eyal G, Aardse R, Lodder BN, Goriounova NA, Asamoah B, B Brakspear AB, Groot C, van der Sluis S, Testa-Silva G, Obermayer J, Boudewijns ZS, Narayanan RT, Baayen JC, Segev I, Mansvelder HD, de Kock CP (2015) Dendritic and axonal architecture of individual pyramidal neurons across layers of adult human neocortex. Cerebral Cortex. 25:4839.

Palmer L, Murayama M, Larkum M (2012) Inhibitory regulation of dendritic activity in vivo. Frontiers in Neural Circuits. 6:26.

Paulk AC, Zhou Y, Stratton P, Liu L, van Swinderen B (2013) Multichannel brain recordings in behaving Drosophila reveal oscillatory activity and local coherence in response to sensory stimulation and circuit activation. Journal of Neurophysiology 110:1703.

Roskies A (1999) The binding problem. Neuron. 24:111.

Sejnowski TJ Churchland PS, Movshon JA (2014) Putting big data to good use in neuroscience. Nature Neuroscience 17:1440.

Sjourn J, Plaais PY, Aso Y, Siwanowicz I, Trannoy S, Thoma V, Tedjakumala SR, Rubin GM, Tchnio P, Ito K, Isabel G, Tanimoto H, Preat T (2011) Mushroom body efferent neurons responsible for aversive olfactory memory retrieval in Drosophila. Nature Neuroscience. 14:903.

Sheffield ME, Dombeck DA (2015) Calcium transient prevalence across the dendritic arbour predicts place field properties. Nature. 517:200.

Stopfer M, Bhagavan S, Smith BH, Laurent G (1997). Impaired odour discrimination on desynchronization of odour-encoding neural assemblies. Nature. 390:70.

Suh GS, Wong AM, Hergarden AC, Wang JW, Simon AF, Benzer S, Axel R, Anderson DJ (2004) A single population of olfactory sensory neurons mediates an innate avoidance behaviour in Drosophila. Nature. 431:854.

Tanaka NK, Tanimoto H, Ito K (2008) Neuronal assemblies of the Drosophila mushroom body. Journal of Comparative Neurology. 508:711.

Tanaka NK, Ito K, Stopfer M (2009) Odor-evoked neural oscillations in Drosophila are mediated by widely branching interneurons. Journal of Neuroscience. 29:8595.

Tang CM, Thompson SM (2012) Perturbations of dendritic excitability in epilepsy. In: Noebels JL, Avoli M, Rogawski MA, Olsen RW, Delgado-Escueta AV editors. Jasper’s basic mechanisms of the epilepsies. 4th ed. (Bethesda (MD): National Center for Biotechnology Information (US) p.494.

Tse D, Langston RF, Kakeyama M, Bethus I, Spooner PA, Wood ER, Witter MP, Morris RG (2007) Schemas and memory consolidation. Science. 316:76.

Palmer LM, Shai AS, Reeve JE, Anderson HL, Paulsen O, Larkum ME (2014) NMDA spikes enhance action potential generation during sensory input. Nature Neuroscience. 17:383.

Vadakkan KI (2007) Semblance of activity at the shared post-synapses and extracellular matrices - A structure function hypothesis of memory (iUniverse Publishers).

Vadakkan KI (2010) Framework of consciousness from semblance of activity at functionally LINKed postsynaptic membranes. Frontiers in Psychology. 1:168.

Vadakkan KI (2011a) A possible mechanism of transfer of memories from the hippocampus to the cortex. Medical Hypotheses. 77:234.

Vadakkan KI (2011b) Processing semblances induced through inter-postsynaptic functional LINKs presumed biological parallels of K-lines proposed for building artificial intelligence. Frontiers in Neuroengineering. 4:8.

Vadakkan KI (2012a) A structure-function mechanism for schizophrenia. Frontiers in Psychiatry. 3:108.

Vadakkan KI (2012b) The nature of “internal sensations” of higher brain functions may be derived from the design rules for artificial machines that can produce them. Journal of Biological Engineering. 6:21.

Vadakkan KI (2013) A supplementary circuit rule-set for the neuronal wiring. Frontiers in Human Neuroscience. 7:170.

Vadakkan KI (2015a) A pressure-reversible cellular mechanism of general anesthetics capable of altering a possible mechanism for consciousness. Springerplus. 4:485.

Vadakkan KI (2015b) A framework for the first-person internal sensation of visual perception in mammals and a comparative circuitry for olfactory perception in Drosophila. Springerplus. 4:833.

Vadakkan KI (2016a) The functional role of all postsynaptic potentials examined from a first-person frame of reference. Reviews in the Neurosciences. 27:159.

Vadakkan KI (2016b) Rapid chain generation of interpostsynaptic functional LINKs can trigger seizure generation: Evidence for potential interconnections from pathology to behavior. Epilepsy and Behavior. 59:28.

Vadakkan KI (2016c) Substantive nature of sleep in updating the temporal conditions necessary for inducing units of internal sensations. Sleep Science. 9:60.

Vadakkan KI (2016d) Neurodegenerative disorders share common features of “loss of function” states of a proposed mechanism of nervous system functions. Biomedicine and Pharmacotherapy. 83:412.

Vadakkan KI (2016e) Examination of memory from a first-person frame of reference provides evidence for a relationship between learning and LTP induction. Biorxiv 10.1101/085589.

Ventura R, Harris KM (1999) Three-dimensional relationships between hippocampal synapses and astrocytes. Journal of Neuroscience. 19:6897.

von der Malsburg C (1999) The what and why of binding: the modeler’s perspective. Neuron. 24:95.

Waddell S (2010) Dopamine reveals neural circuit mechanisms of fly memory. Trends in Neurosciences. 33:457.

Wilson CJ (1984) Passive cable properties of dendritic spines and spiny neurons. Journal of Neuroscience. 4:281.

Wise RA (2004) Dopamine learning and motivation. Nature Review Neuroscience. 5:483.

Wong AM, Wang JW, Axel R (2002) Spatial representation of the glomerular map in the Drosophila protocerebrum. Cell. 109:229.

Xie L, Kang H, Xu Q, Chen MJ, Liao Y, Thiyagarajan M, O’Donnell J, Christensen DJ, Nicholson C, Iliff JJ, Takano T, Deane R, Nedergaard M (2013) Sleep drives metabolite clearance from the adult brain. Science. 342:373.

Yagishita S, Hayashi-Takagi A, Ellis-Davies GC, Urakubo H, Ishii S, Kasai H (2014) A critical time window for dopamine actions on the structural plasticity of dendritic spines. Science. 345:1616.

Yasuyama K, Meinertzhagen IA, Schrmann FW (2002) Synaptic organization of the mushroom body calyx in Drosophila melanogaster. Journal of Comparative Neurology. 445:211.

Zeron MM, Hansson O, Chen N, Wellington CL, Leavitt BR, Brundin P, Hayden MR, Raymond LA (2002) Increased sensitivity to N-methyl-D-aspartate receptor-mediated excitotoxicity in a mouse model of Huntington’s disease. Neuron. 33:849.

## References

Abbott LF (2008) Theoretical neuroscience rising. Neuron. 60:489.

Dudek SM, Alexander GM, Farris S (2016) Rediscovering area CA2: unique properties and functions. Nature Review Neuroscience. 17:89.

Hatanpaa KJ, Raisanen JM, Herndon E, Burns DK, Foong C, Habib AA, White CL 3rd. (2014). Hippocampal sclerosis in dementia epilepsy and ischemic injury: differential vulnerability of hippocampal subfields. Journal of Neuropathology and Experimental Neurology. 73:136.

Johnston D, Brown TH (1981) Giant synaptic potential hypothesis for epileptiform activity. Science. 211: 294.

Lledo PM, Zhang X, Sdhof TC, Malenka RC, Nicoll RA (1998) Postsynaptic membrane fusion and long-term potentiation. Science. 279:399.

Michael BD, Solomon T (2012) Seizures and encephalitis: clinical features management and potential pathophysiologic mechanisms. Epilepsia. 53 Suppl 4:63.

Sahin M, Sur M (2015) Genes circuits and precision therapies for autism and related neurodevelopmental disorders. Science. 350 (6263) pii: aab3897.

Segal IS, Walton JK, Irwin I, DeLanney LE, Ricaurte GA, Langston JW, Maze M (1990) Modulating role of dopamine on anesthetic requirements. European Journal of Pharmacology. 186:9.

Singh SP, He X, McNamara JO, Danzer SC (2013) Morphological changes among hippocampal dentate granule cells exposed to early kindling-epileptogenesis. Hippocampus. 23:1309.

